# Atlantis: An integrative database for human proteome structural and functional sites

**DOI:** 10.64898/2026.08.29.747983

**Authors:** Natalia De Oliveira Rosa, Piergiorgio Ferronato, Martina Varisco, Marin Matic, Mirko Ruscio, Pasquale Miglionico, Francesco Raimondi

**Affiliations:** Laboratorio di Biologia Bio@SNS, Scuola Normale Superiore, Piazza dei Cavalieri 7, 56126, Pisa, Italy

## Abstract

Understanding protein mechanisms in health and disease requires characterizing the functional roles of individual amino acid residues. To explore the role of residues and their mutations, we have developed Atlantis, a database that integrates structural and functional information at the human proteome residue level. A graph database enables complex queries and the retrieval of integrated information for multiple functional analysis of protein systems. A Model Context Protocol (MCP) connector allows the interrogation of the resource through Large Language Models (LLMs) or agentic frameworks for biomedical research.

Atlantis annotates over 11M residues across 20k human proteins, identifying hundreds thousands intra- and inter-protein contacts in PDB as well as AlphaFoldDB structures. We also provide the possibility to analyze and integrate predicted 3D complexes inputted by the user, and we showcased these features on hundreds of AlphaFold-multimer complexes of GPCRs and LRRK2 interaction networks. The tool is freely accessible at https://atlantis.bioinfolab.sns.it/.

**GRAPHICAL ABSTRACT:** 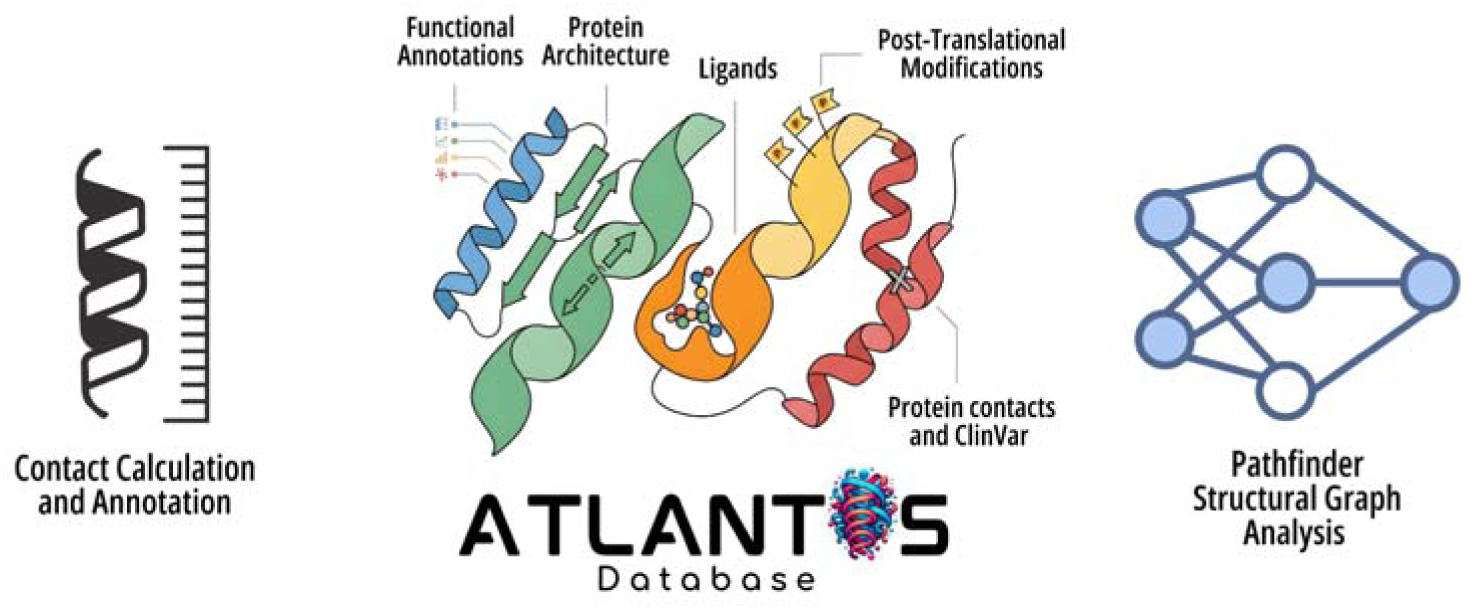

## INTRODUCTION

Characterizing the functional roles of individual amino acid residues is fundamental to deciphering protein mechanisms and their implications for health and disease. There are several established protein knowledge bases. For instance, UniProtKB^1^ offers comprehensive sequence annotations and functional information, including variant data and links to structures. More structure-centered web servers, such as SWISS-MODEL^2^ and AlphaFold DB^3,4^, also provide the option to visualize some sequence annotations (i.e., domains, pathogenicity) on the structures. These tools have transformed the work of both experimental and computational biologists, allowing them to easily access information that would otherwise require a very time-consuming literature mining. However, they often do not provide an easy and user-friendly way to access the data in the residue-centric fashion necessary for detailed functional analysis. Similarly, while other valuable resources like the Protein Contact Atlas^5^, AlphaSync^6^ or Residue Interaction Network Generator (RING)^7^ compute detailed protein contacts and geometric descriptors, they generally lack a straightforward way to map this structural information back to UniProt sequences and annotations for integrated analysis. Therefore, even if many tools are available for the prediction of the functional impact and/or pathogenicity of human variants^8,9^, it is still difficult to mechanistically interpret the likelihood scores provided by the models, and few tools (e.g., Mechismo^10^ or EXPANSION^11^) have attempted to tackle this issue.

Over the years, several dedicated databases have also been developed to capture the complex relationships between post-translational modifications (PTMs), structural features, and disease-associated variants. For instance, tools such as PTMcode^12^ and PRISMOID^13^ allowed the exploration of functional associations and crosstalk between PTMs within proteins, but they are not anymore maintained and accessible. Similarly, Missense3D- PTMdb^14^ leverages AlphaFold models to visualize the structural impact of missense variants and PTMs, but the amount of information it integrates is limited to these two features. While these resources offer invaluable insights into PTM and variant mapping, combining these functional annotations with residue-residue contact networks could provide a more complete, mechanistic understanding of protein allostery and signal propagation. This approach has proven highly effective in identifying structural hubs and communication pathways^15^. Tools like webPSN^16,17^ and Network Analysis of Protein Structures (NAPS) server^18^ have facilitated the investigation of structural communication and allosterism, but lack an automatic cross-referencing to functional annotation.

To address this gap, we present Atlantis, a novel database offering an integrative view of structural and functional sites within the human proteome at the residue level.

Atlantis uniquely provides access to pre-calculated intra- and inter-protein contacts derived from both experimental structures (PDB)^19^ and AlphaFold-predicted models (AFDB)^3^, alongside curated data from diverse sources, including sequence annotations, protein structures, protein and ligand interactions, ClinVar^20^ annotated mutations and pre-computed variant-effect prediction^8,9^. This integrated approach facilitates the visualization of functional annotations and variants on 3D structures, enabling insights into how amino acid changes drive alterations at the structural and functional level (Figure 1).

**Figure 1.**
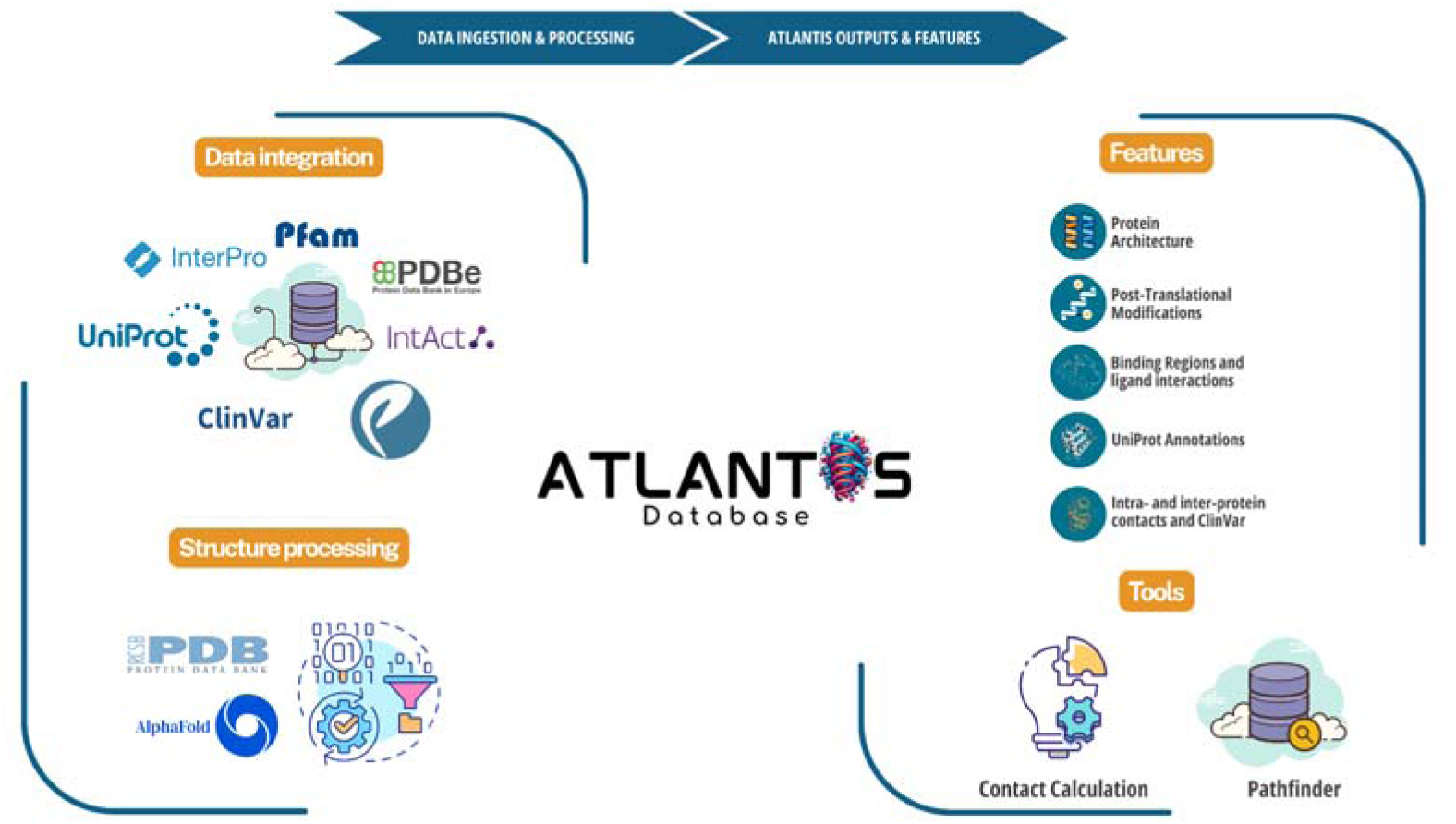
Overview of the ATLANTIS database workflow and architecture. The pipeline is divided into two main phases: data ingestion and processing (left) and user outputs and features (right). First, functional and structural data are retrieved from various external repositories (Pfam, InterPro, UniProt, IntAct, ClinVar, PDB, and AlphaFold). These data undergo rigorous integration and structure processing pipelines to generate the ATLANTIS database. On the user end, ATLANTIS provides comprehensive structural annotations (Features), including protein architecture, post-translational modifications, binding regions, and intra-/inter-protein contacts. Furthermore, the platform offers dedicated computational “Tools” such as the Pathfinder algorithm for structural graph analysis and custom Contact Calculation modules.

Moreover, Atlantis introduces the Pathfinder module to perform integrative structure contact network analysis. Pathfinder utilizes a graph architecture to investigate protein structures as contact networks, enriching topological analyses (like shortest paths and centrality metrics) with cross-referencing of variants, ligands, and PTMs. Atlantis empowers researchers to programmatically retrieve residue-level functional annotations at a proteome-wide scale, as well as investigate the roles of specific residues in biological processes through a user-friendly web interface and GraphQL API as well as through Large Language Models (LLMs) through a Model Context Protocol (MCP) connector. We also provide the possibility to process experimental or predicted 3D structures inputted by the users.

By providing an advanced, residue-centric resource, Atlantis represents a significant step forward in unraveling the complexities of protein function and molecular interactions.

## METHODS AND DESCRIPTION

### Data collection and functional annotation

We retrieved residue-level annotations from UniProtKB (Release v2026_01)^1^, including: molecule processing, membrane topology, secondary structure elements, natural variants, binding pockets, active sites, disulfide bridges, and crosslinks. To maintain high-fidelity annotations, the Atlantis database undergoes synchronized biannual updates across all variant and functional annotation sources

Protein architecture was defined by integrating domain and family definitions retrieved from UniProtKB^1^, InterPro (v108.0)^21^, and Pfam (v38.1)^22^. To ensure residue-level correspondence, protein sequences were aligned against the Pfam-A HMM library using hmmalign^23^ from the HMMER suite (v3.3.2)^24^. To optimize the visual representation of InterPro annotations in the web interface, a hierarchical rendering strategy based on category relevance (Domain > Homologous Superfamily > Repeat > Family) was implemented. In cases of spatial conflict within the viewer, a coverage-based logic is applied: overlapping features are merged if the overlap exceeds 70% of their length, whereas only the longest annotation is retained for intermediate overlaps (30–70%).

Protein–protein interaction binding regions were derived from the IntAct database (accessed v2026-01-14)^25^ by extracting experimentally verified residue ranges from PSI-MI TAB files. High-confidence PTMs, encompassing modification types and site positions, were obtained from PhosphoSitePlus® (Release 04/2025)^26^.

For the structural analysis, we collect all the available human 3D structures from PDB (experimental) and the AlphaFold Protein Structure Database (v6)^3,4^. For AF-predicted structures, Local Distance Difference Test (pLDDT) scores and secondary structure assignments were extracted directly from AlphaFold mmCIF files. Besides structures from public databases, we also included thousands of AF-multimer-predicted complexes, including heterotrimeric G proteins in complex with GPCRs ^27,28^ and Adenylate Cyclase (AC), as well as a recently computed LRRK2’s interactome^29^.

### Structural processing and contact definition

Residue-residue contacts were defined using a Cβ distance cutoff of 8 Å (using Cα for Glycine)^30^. Contacts were computed using an in-house pipeline implemented with cifparse-obj^27^. Residues from experimental structures were mapped to UniProtKB reference sequences via SIFTS XML integration^31^.

### Prediction of AC-Gα complexes via AlphaFold-Multimer

Structural predictions of six Gαs (GNAI1, GNAI2, GNAI3, GNAO, GNAS, GNAZ) in complex with full length Adenylate Cyclase (ADCY1-9) were performed with AlphaFold2-multimer (v2.3)^32^ using the ColabFold (v1.5.2) implementation^33^. For each of the 54 complexes, 5 distinct models were generated using 5 recycles. All predictions utilized the default multimer model type to predict the heterodimeric assemblies from their respective FASTA sequences retrieved from UniProt. Among the 5 models generated for each AC-Gα pair, the one with the highest confidence score was relaxed using Rosetta (v2021.16+release.8ee4f02)^34^.

### Ligand integration and binding site mapping

Ligand metadata, specifically physicochemical classifications (e.g., cofactor-like, drug-like, reactant-like, ion, small molecule), were retrieved from PDBe (accessed June 30, 2026)^31,35^, while external cross-references were derived directly from ChEMBL^36^. Furthermore, ligand-binding sites within experimental PDB structures were spatially defined using a dedicated structural processing pipeline^27^. Residues were classified as part of the binding interface if the distance between any ligand atom and the residue Cβ atom (or Cα for Glycine) was less than 8 Å. For each interaction, the specific atom pair corresponding to the minimum distance was retained.

### ClinVar variants and pathogenicity scoring

Disease-associated genetic variation was incorporated by retrieving human variants from the ClinVar database^20^. These variants were systematically mapped to their corresponding protein sequence residues. To provide a prediction of the variant impact, we annotated the dataset with AlphaMissense pathogenicity scores, using the Ensembl Variant Effect Predictor (VEP)^37^.

### Graph-based topological modeling

To enable integration of the different layers of structural and functional annotations, the datasets, initially consolidated in a relational MySQL environment, were transformed into a graph architecture using Neo4j. The graph schema was designed to mirror biological hierarchy while encoding 3D spatial relationships (i.e. 3D contacts) as traversable edges. Subsequently, the graph was enriched by mapping functional features to their corresponding node.

### Pathfinder Module

The Atlantis Pathfinder module utilizes a Neo4j-powered Knowledge Graph to analyze protein structures as networks in which nodes are protein residues and edges are contacts (defined as described in “Structural processing and contact definition”). Each analytical level is designed to answer a specific biological question about a protein: (i) how structural information propagates through a fold, (ii) how post-translational modifications are organized and regulated, and (iii) how genetic variants relate to structure and function.

### Structural analyses

This tier addresses how a protein’s 3D geometry is linked to function: which minimal residues connect two functional sites, which residues are structurally indispensable, how the assembly of a complex can be perturbed, and how a structure’s contact architecture changes across conformational or predicted states.

#### Level 1: Physical Pathways

The tool identifies the shortest path between two residues or ligands in the contact network computed with the Dijkstra’s algorithm^38^ via the Neo4j Graph Data Science (GDS) library (gds.shortestPath.dijkstra.stream), using distances as edge weights under a user-selected Path Weighting scheme (Geometric vs. Functional). For each node of the identified path, a cross-reference query retrieves functional annotations.

#### Level 2: Structural Hubs

Calculates betweenness centrality and degree^39^ via the Neo4j GDS library on intra-chain contacts, inter-chain and ligand contact.

#### Level 3: Structure differences (PDB vs AlphaFold)

Evaluates local similarity between a user-defined PDB and its corresponding AlphaFold model by comparing, for each corresponding residue, both the change in contact degree and the overlap (Jaccard index) between the two networks’ contact-partner sets, supported by automated 3D superposition in the integrated Mol* viewer.

#### Level 4: Conformational Differences (PDB vs PDB)

Evaluates local similarity between two experimental structures of the same protein by comparing, for each residue resolved in both, the change in contact degree and the overlap (Jaccard index) between the two networks’ contact-partner sets, supported by automated 3D superposition.

### Post-translational modification analyses

This tier addresses the spatial organization of post-translational modifications, and its implications for regulatory mechanisms.

#### Level 5: PTM Crosstalk (3D Regulation)

Maps PTMs onto the contact network to identify spatially close post-translationally modified residues, classifying pairs as same-residue, proximal, or distal based on sequence separation, and detecting higher-order motifs (trios and tetrads) as fully connected cliques within the contact graph.

#### Level 6: PTM Hotspots

Employs a 31-residue sliding window algorithm, evaluated against a 1000-fold permutation null with a permutation-based FDR correction^40^, to identify sequence regions exhibiting statistically significant PTM enrichment.

#### Level 7: Domain-PTM Enrichment

Maps PTMs onto functional domains annotated by UniProt, Pfam, and InterPro. By calculating the ratio of modified residues relative to the total domain length, the module systematically ranks functional domains based on their PTM density.

#### Level 8: PTM Structural Enrichment Analysis

Identifies mutations that directly affect PTM residues, as well as mutations affecting residues in contact with a PTM site. A two-sided Fisher’s exact test then evaluates whether the pathogenic-versus-benign composition differs significantly between variants near PTM sites and those farther away.

### Variant analyses

This tier addresses the structural and evolutionary context of individual variants: their relationship to complex assembly and ligand binding, their conservation across homologous proteins, their spatial co-occurrence with other pathogenic variants, and inference of effect based on structural evidence.

#### Level 9: PPI Interface Hotspots

Systematically cross-references inter-chain contacts on a given structure with ClinVar variants to identify mutations that may affect quaternary assembly, and tests by Fisher’s exact test whether pathogenic variants are enriched at the interface, prioritising each site by the pathogenic density of its 3D neighbourhood and the number of partners it contacts.

#### Level 10: Functional Sites Overlap

Intersects UniProt-annotated ligand-binding and active sites with ClinVar variants to identify mutations positioned to disrupt ligand or cofactor binding or catalysis, either directly or through a residue in 3D contact (≤8 Å), assessing a variant’s functional consequence at a defined functional site. A Fisher’s exact test assesses whether pathogenic variants are enriched at these sites relative to benign ones, and a cluster-synergy analysis flags spatial groupings in which several variants converge on the same site.

#### Level 11: Evolutionary Mapping (HMM)

Identifies conserved sites across protein families^41^ enriched in mutations. To avoid the computational burden of on-the-fly alignments, protein sequences are pre-aligned against the Pfam-A HMM library using hmmalign, so that each residue of a query protein is mapped to the consensus column of its Pfam domain family. This allows a pathogenic ClinVar variant on one human paralog to be instantly cross-referenced onto the corresponding consensus position of any other human protein with the same domain, surfacing candidate vulnerability at that position.

#### Level 12: Structural Co-Pathogenicity

Identifies pairs of amino acids that are interacting (≤8 Å) where each one has a pathogenic mutation in the ClinVar database, suggesting that the disease affects two residues involved in an interaction. The inter-contact residue pairs identify pathogenic residue pairs in protein-protein interactions, while intra-contact residue pairs identify residues close to each other with pathogenic mutations.

#### Level 13: VUS Structural Prioritisation

Lists residues in 3D contact with a target variant of uncertain significance (VUS) to identify direct spatial proximity to established ClinVar pathogenic variants, catalytic active sites, or ligand-binding pockets, allowing VUS reclassification based on the structural context.

#### Level 14: Variant Convergence Triage

When provided with a list of variants by the user, typically a list of VUS, this level combines the data gathered at structural, functional, and evolutionary levels above and provides a per-residue count of lines of evidence that are active without making any kind of prediction score out of their combination. It prioritizes variants neighboring functional sites such as those involved in ligand binding or catalysis, PTMs, and conserved across families. This enables quick prioritization of the variants based on the lines of evidence from structure and evolution.

### Contact Calculation Module

The Contact Calculation module integrates a pipeline for the coordinate processing and functional annotation of user-provided 3D structures (via mmCIF upload, PDB identifiers or UniProt accessions to retrieve predicted structures from AlphaFoldDB). The input structure is parsed as described in “Structural processing and contact definition” to build a contact network using a customizable residue distance threshold (default: 8 Å). The generated contact network is analyzed using NetworkX. By utilizing physical distances as edge weights, the backend computes shortest paths between residues via Dijkstra’s algorithm and identifies critical structural hubs using betweenness centrality. All analytical tasks are secured by a unique UUID to ensure session persistence and data reproducibility.

To establish biological identity, the pipeline extracts protein sequences from the structural data and executes a BLASTP search against a local UniProtKB/Swiss-Prot database. This sequence alignment synchronizes the input coordinates with Atlantis’s functional repository.

### Web-server implementation

ATLANTIS is freely accessible at https://atlantis.bioinfolab.sns.it. The frontend interface was developed using ReactJS (v18.2.0), leveraging PrimeReact (v10.5.0) and React-Bootstrap (v2.10.0) for UI components. High-performance interactive 3D visualisation is supported by the Molstar viewer (v4.1.0). The backend infrastructure is implemented in Python, utilising FastAPI (v0.104.1) for HTTP request management and Strawberry GraphQL (v0.217.1) for API query orchestration. Data persistence relies on a hybrid architecture comprising MySQL for structured annotation retrieval and Neo4j (v4.1.13) for graph-based operations. Computational routines for network topology analysis were performed using NetworkX (v3.4.2), while structure parsing and contact calculations were handled by Biopython (v1.83) and the optimised cifparse-obj module (v7.105), respectively. Sequence mapping to UniProt accessions is facilitated by BLASTP (v2.10.1+).

### API

Direct programmatic access to the Atlantis dataset is provided via an Application Programming Interface (API) to facilitate high-throughput analysis. The interface was implemented in Python, leveraging FastAPI (v0.104.1) and the Strawberry-GraphQL framework (v0.217.1). Through the primary endpoint (https://api.atlantis.bioinfolab.sns.it/graphql), specific data layers—encompassing residue-level annotations, intra- and inter-chain structural contacts, ligand interaction profiles, and ClinVar variants can be retrieved. Comprehensive documentation, including schema definitions and syntax guidance, is available at https://atlantis.bioinfolab.sns.it/api/ to support integration into external computational pipelines.

### Model Context Protocol (MCP) server

To further extend accessibility for AI-assisted analysis, the Atlantis platform additionally provides a Model Context Protocol (MCP) server, allowing large language model (LLM)- based assistants to query the underlying dataset through natural language rather than direct GraphQL syntax. The server was implemented using the FastMCP framework (v3.4.7) and is accessible at the endpoint https://mcp.atlantis.bioinfolab.sns.it/mcp. Documentation describing client configuration and representative example queries is available at https://atlantis.bioinfolab.sns.it/mcp/, enabling retrieval of residue-level annotations, intra- and inter-chain structural contacts, ligand interaction profiles, and ClinVar variant data through conversational or agentic interfaces integrated into external analytical workflows.

## RESULTS

### Using the WebServer

The main interface supports precise queries via gene symbols, UniProt accessions, or specific residues. An advanced query module allows multiple filtering schemes to generate custom datasets. The “Tools” menu in the global navigation bar hosts the Contact Calculation module for the analysis of user-provided structures and Pathfinder module for the exploration of structure-specific residue-residue contact networks. The platform includes dedicated sections for methodology (“About”), comprehensive usage instructions (“Help”, “API” and “MCP”), and a “Release Log” documenting database versioning and the latest update cycles, complemented by a frequently asked questions (“FAQs”) repository and glossary. To facilitate external research, all modules include export/download options and persistent URL sharing for direct collaborative access to specific filtered outcomes.

### Interactive data integration and multi-dimensional analysis

Upon executing a query, the Results Page serves as an integrated analytical dashboard, organised into six specialised tabs: Overview, Functional Site Information, Intra-protein Contacts, Inter-protein Contacts, Ligands, and ClinVar. The Overview shows an interactive dashboard of feature counts that function as direct hyperlinks to granular data. Detailed residue-level insights are centralised in the Functional Site Information section, featuring a 23-column comprehensive annotation table and an interactive Feature Viewer for linear mapping of functional annotations. Annotations are projected onto the AlphaFold 3D structural models; users can perform PDB superposition to align experimental structures with predicted models, enabling direct comparison of functional site coordinates across structural states. This section further incorporates structures from AlphaFill^42^, allowing users to “transplant” ligands or metal ions into models to enhance functional interpretations. For residues involved in protein-protein interactions, the Intact binding region visualization is complemented by an interactive Network viewer, which visualises the protein’s interactome. Contact and Ligand tabs use a Mol* 3D viewer to visualise residue-residue and residue-ligand contacts. In the Contact tabs, the analysis is supplemented by contact heatmaps and statistical distribution plots, where users can select table rows to highlight specific contacts in the 3D structure. In the Ligands tab, the structure section provides a list of bound molecules; users can select a specific ligand to highlight its binding pocket in the Mol* view and visualise its neighbour residues. Lastly, the ClinVar tab offers a specialised viewing environment where variants are mapped onto 3D structures. This section is enhanced by a Lollipop plot interface that allows the selection of variants by clinical significance or AlphaMissense class. Both the Contact and ClinVar tabs integrate also new AF-multimer models, including complexes of G proteins with GPCRs and Adenylate Cyclase (AC), as well as the LRRK2 interactome, to improve the contextualization of interactions and disease variants within physiological macromolecular assemblies. Beyond monomeric structures, Atlantis also integrates 7,686 predicted quaternary structures from the AlphaFold Database, comprising 1,624 high-confidence homodimeric and 6,062 heterodimeric protein complexes predicted at proteome scale. These multimeric assemblies are directly accessible to users within the Functional Sites, Contacts, ClinVar Variants, and Pathfinder modules, allowing structural and variant analyses to be interpreted in the context of the relevant protein complex rather than the isolated chain alone.

### Advanced Search

The Advanced Search interface allows users to construct custom datasets using advanced filtering criteria. Generated results are presented in a sortable table that provides a comprehensive overview of matching proteins or specific residues. Key metadata, including curation status, gene symbols, UniProt accessions, and sequence length, are displayed alongside a Protein Features Overview, which quantifies the density of domains, structural elements, and interaction sites for each entry. Designed as an interactive hub, this page allows users to track applied filters and navigate from the summary list to detailed, protein/residue-specific annotation pages by clicking on the respective gene or residue identifiers.

### Residue Tag System

Atlantis implements a Residue Tag System that condenses complex multidimensional data into an “at-a-glance” iconic summary located within the primary results table. Distinct interactive icons flag critical properties, including experimentally verified and AlphaFold-predicted contacts (intra- and inter-chain), ligand binding events, and the presence of ClinVar pathogenicity data. Additionally, a specific “Functional Significance” tag aggregates diverse annotations, ranging from PTMs and active sites to domain membership, allowing users to instantly identify and prioritise biologically relevant residues without navigating through multiple sub-sections.

### Contact Calculation Module

The Contact Calculation Module allows users to process custom structural models to identify contacts and map functional hotspots. Once a structure is submitted, the system performs an automated cross-reference, projecting Atlantis’s repository of functional annotations (such as PTMs, clinical variants, and domains) directly onto the user-provided model. These mapped features are instantly visualized through the Residue Tag System, enabling an “at- a-glance” functional assessment of the novel structure’s topology (Figure 2A). Structural contacts can be viewed in the interactive 3D environment (Figure 2B) or exported in tabular format.

**Figure 2.**
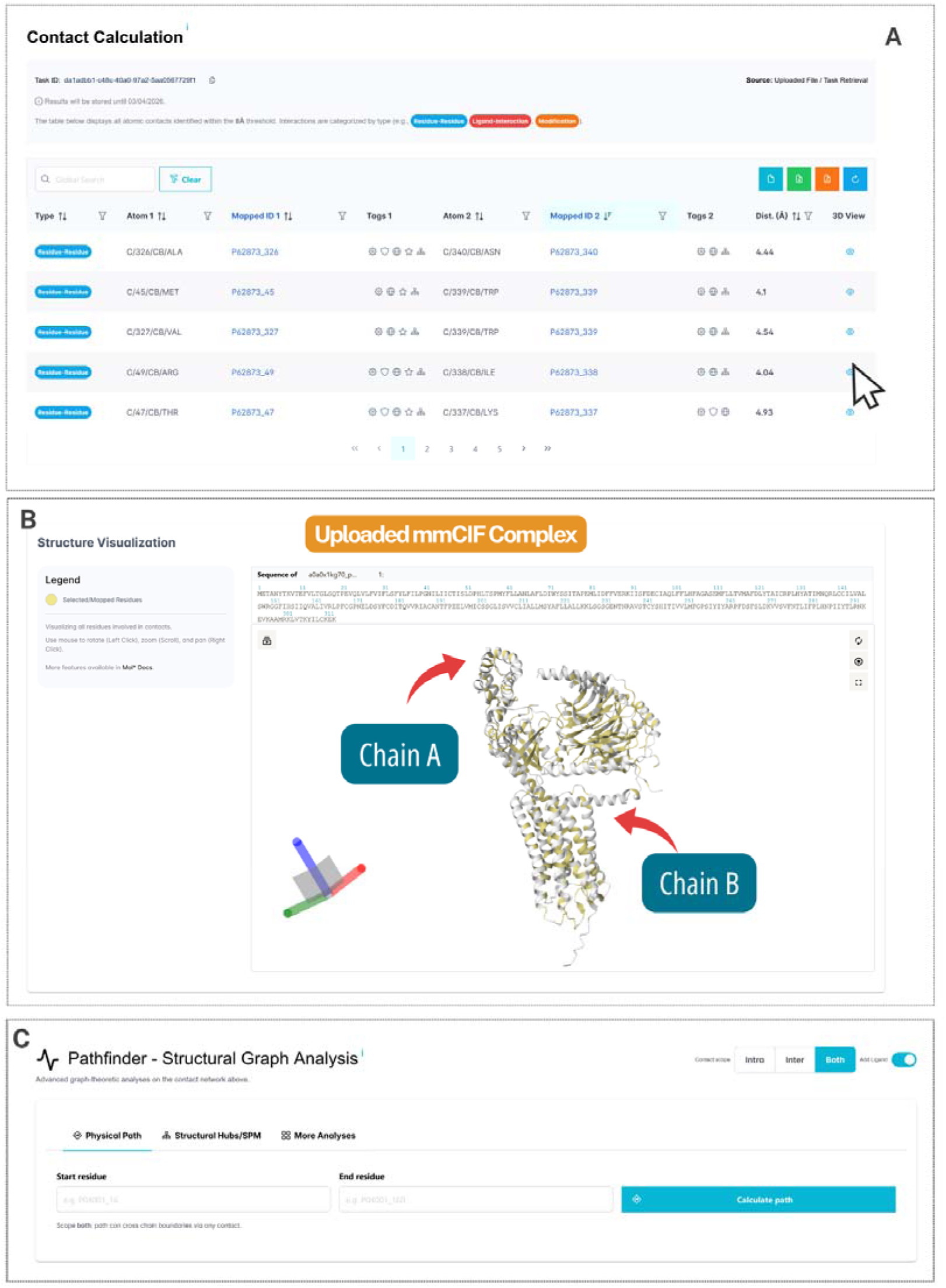
Overview of the ATLANTIS web server interface and Contact Calculation Module. **(A)** Contact Calculation results table displaying inter- and intra-protein contacts alongside the Residue Tag System. A dedicated “3D View” button renders specific interactions. **(B)** Interactive structure visualization (Mol* viewer) mapping calculated contacts onto macromolecular complexes. **(C)** Downstream Graph Analysis interface allowing to isolate shortest communication pathways and Centrality measures to identify critical Structural Hubs.

The analytical workflow further extends into a Downstream Graph Analysis panel (Figure 2C). Here, users can specify origin and target residues to compute the shortest communication path, generating a trajectory report that details the accumulated distance and functional profile of each intermediate node. Moreover, the module creates a custom metrics table to rank Structural Hubs according to their centrality and thus helps in focusing on the important nodes that act as intermediaries to facilitate network communication for further study. The software has a More Analyses (Functional Annotation) tab, which includes 3D distances along with the sequence annotation features like ClinVar variants, PTMs, and ligand binding sites using features like Ligand-Variant Overlap, PTM Crosstalk 3D, PTM Hotspots, PTM-Variant Disruption, and the VUS Resolver.

### Pathfinder Module

Through the Pathfinder interface, users can execute topological and functional analyses on protein structures (see Methods). The input varies according to the selected level, accepting HGNC gene symbols, UniProt accessions, PDB IDs, or AlphaFold models, alongside specific sequence positions. Upon execution, the backend Neo4j graph is interrogated on the fly, returning results via an interactive dashboard. To facilitate structural interpretation, the interface natively integrates the Mol* viewer for 3D inspection of shortest paths, centrality hubs, and mutation microenvironments. Additionally, for sequence-level statistical clustering, interactive density plots are rendered using Plotly. Furthermore, all extracted topological metadata and cross-referenced ClinVar annotations are organized into responsive DataTables, which can be filtered and directly downloaded in CSV format.

### Pathfinder recovers established allosteric, regulatory, and disease mechanisms across three protein systems

We validated Pathfinder’s structural graph analyses against three proteins with experimentally characterized regulatory architectures: for the structural tier, the β2- adrenergic receptor (ADRB2), a class A GPCR with a mutagenesis- and MD-defined activation pathway; for variants, transthyretin (TTR), a homotetramer with specific interface variants causing amyloidosis; for PTMs, the tyrosine kinase c-Src, with a well-defined post-translational autoinhibitory mechanism.

### ADRB2: structural changes for signal transduction

ADRB2 has well-characterized signal propagation routes from the orthosteric pocket across the transmembrane bundle up to the intracellular G-protein-coupling surface, and we tested if Pathfinder could reproduce it only based on the graph. Using Level 1 (Physical Pathways), which computes the Dijkstra shortest weighted path between two user-defined residues, we queried the agonist-bound, Gs-coupled structure (PDB 3SN6 ^43^) between the orthosteric anchor Asp113 and the DRY arginine Arg131 in the G protein cavity. The returned path threaded directly through the CWxP toggle (Cys285/Trp286) and the PIF connector (Phe282) before reaching the TM5 pivot Tyr219 (55.93 Å) (Figure 3A), reproducing the activation cascade defined by mutagenesis and molecular dynamics ^44,45^. Repeating the same query and endpoints on a fusion-free inactive structure (PDB 2R4R/2R4S ^46^) returned a different route, rerouting around the periphery (TM2–TM3, 48.0 Å) and touching none of these switches (Figure 3B). A second, independent Level 1 query on the sodium- pocket/NPxxY coupling (Asp79→Tyr326) showed the same pattern: direct engagement through Asn322 in the NPxxY motif when active (17.55 Å), a detour through Asn51 when inactive (21.3–21.5 Å).

**Figure 3.**
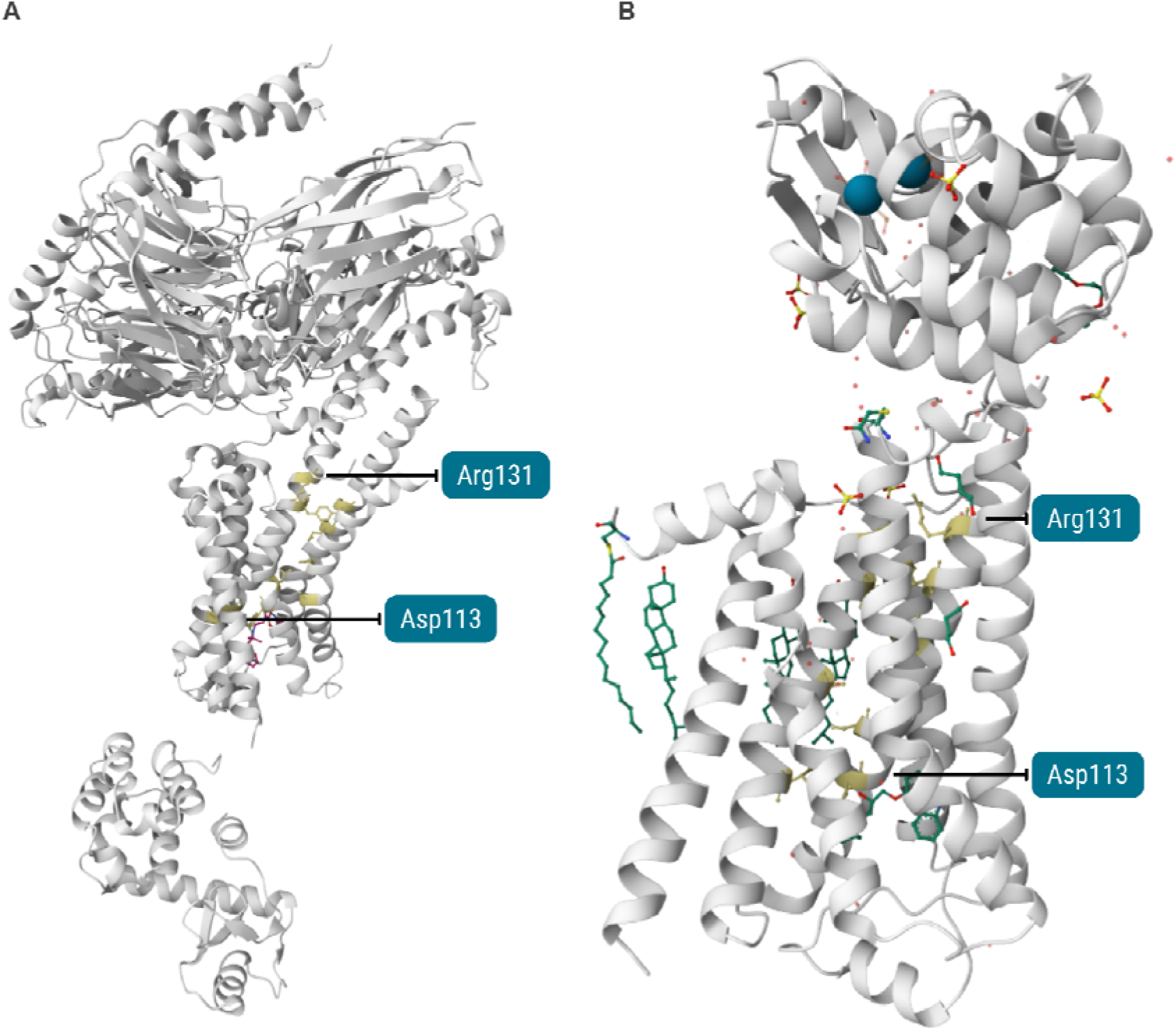
Pathfinder Level 1 (Physical Pathways) shortest weighted paths in ADRB2, rendered in the interactive structure viewer (Mol* viewer) of the Atlantis web interface. Path residues are shown as yellow sticks on white cartoons. Path residues are shown as yellow sticks on white cartoons. **(A)** Active, Gs-coupled receptor (PDB 3SN6), where the 55.93 Å path crosses the CWxP toggle (Cys285/Trp286), the PIF connector (Phe282) and the TM5 pivot Tyr219. **(B)** Inactive receptor (PDB 2R4R/2R4S), where the same query returns a shorter peripheral TM2–TM3 route (48.0 Å) bypassing all activation microswitches.

We then assessed whether structural hubs identified by degree and betweenness centrality were preserved using Level 2 on the full network (intra- and inter-chain contacts, ligand included) and a restricted network (intra-chain only, ligand excluded). The current analysis identified 21 hubs in both networks, with persistent hubs mapping to established structural regions including the sodium pocket and TM1–TM3 packing core, as well as the PIF/CWxP and NPxxY microswitch regions. Their persistence, after ligand and inter-chain contacts were excluded, indicates that these central positions are supported by the receptor’s intrinsic structural network. Structural hubs partially overlapped with residues critical for signal transduction, suggesting a prioritization tier for mutagenesis, with residues retrieved by both Level 1 and Level 2 representing candidates for structural perturbation and residues identified primarily by Level 1 representing candidates more directly associated with signal propagation.

Finally, to evaluate structural changes during activation, we used Level 3 (Structure Differences), to compare the intra-chain contact network of the active PDB structure to that of its corresponding inactive AlphaFold model. The residues with a change in contact degree map closely onto the established activation microswitches. The active, Gs-bound structure carries fewer intramolecular contacts at these positions than the AlphaFold model, consistent with the DRY, PIF-connector, and NPxxY microswitches each shedding a portion of their intramolecular packing as the receptor engages the G protein: Arg131 in the DRY-motif loses 3/6 contacts upon activation, and its ionic-lock partner Glu268 gain 2/1 contacts; two further residues flanking the PIF connector (212, 279) go from 6 to 4/3 contacts. In addition, we identified four hinge residues (i.e., four residues with similar degrees but different contacts): Tyr326 (Y7×53, NPxxY motif, TM7), Leu275 (L6×37, TM6), Arg131 (R3×50, DRY motif, TM3) and Asn103 (3×22, extracellular end of TM3). All residues undergoing structural changes are additional candidates for mutations stabilizing a specific conformation.

### Transthyretin: variants disrupting the interface and ligand pocket, and VUS reclassification

Transthyretin (TTR) is a homotetrameric plasma and cerebrospinal-fluid protein that transports thyroxine and indirectly vitamin A, whose native tetramer can dissociate into misfolding-prone monomers in the presence of specific point mutations, causing hereditary amyloid disease. Here, we validate Pathfinder variant tier against TTR’s established amyloidogenic mutations, testing interface and ligand-pocket variants.

Level 9 (PPI Interface Hotspots), which cross-references inter-chain contacts with ClinVar variants, was applied to the thyroxine-bound tetramer (PDB 2ROX). Ranked by interface promiscuity, only two positions scored above zero: 122 (p.Val142 in precursor numbering), which carries the archetypal cardiac amyloidosis variant Val122Ile (p.Val142Ile) ^47^, and 114 (p.Tyr134) (Figure 4A). It also recovered position 119 (Thr119Met/p.Thr139Met) with a ClinVar tag of conflicting classifications: T119M is in fact a protective, dissociation-slowing variant with contested clinical classification ^48^. Position 114 showed substitution-specific pathogenicity within a single site (Y114H/C/S pathogenic, Y114N likely benign) (Figure 4B).

**Figure 4.**
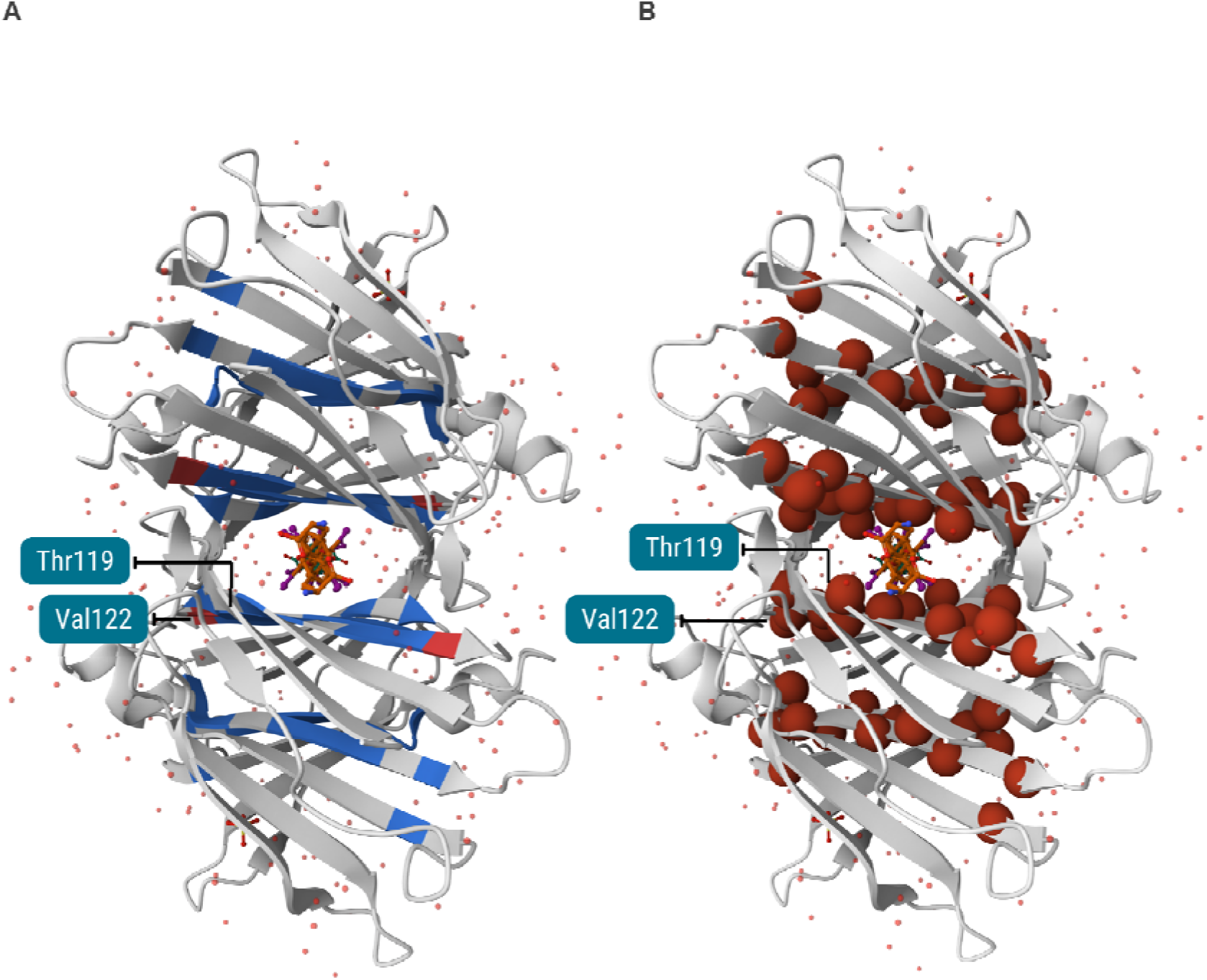
Pathfinder Level 9 (PPI Interface Hotspots) applied to the thyroxine-bound transthyretin tetramer (PDB 2ROX), rendered in the interactive structure viewer (Mol*) of the Atlantis web interface. The bound thyroxine is shown as orange sticks and ordered waters as small red spheres. **(A)** Residues coloured by promiscuity score, which counts how many distinct interface partners each residue contacts: in this structure only two positions score above zero. The top-ranking hotspot Val122 and the conflicting-classification site Thr119 are labelled. **(B)** The same structure with ClinVar variant positions shown as red spheres.

Because the interface hotspots are close in sequence and in 3D structure, we applied Level 10 (Functional Sites Overlap), which intersects UniProt-annotated ligand-binding regions with ClinVar variants, to assess if they are in a ligand-binding pocket. Seven of thirteen flagged positions are canonical thyroxine halogen-binding-pocket residues (Met13, Thr106, Ala108, Ala109, Leu110, Ser117, Glu54; UniProt P02766 precursor positions 33, 126, 128, 129, 130, 137 and 74 respectively), and the analysis additionally flagged Leu55Pro (Leu75Pro in precursor numbering). Level 9 and Level 10 overlap at two positions, Leu110 and Ser117 (precursor 130 and 137), both of which line the thyroxine channel at the dimer– dimer interface. The two modules are otherwise largely complementary: Level 9 additionally highlights the 87–96 and 112–122 strands, whereas Level 10 recovers the 13–24 and 49–55 blocks that Level 9 does not reach.

Thanks to the high ClinVar coverage, we could re-evaluate the pathogenicity of VUS using level 13 (VUS Structural Prioritisation): 88% of 100 VUS showed direct contact with an already-confirmed pathogenic residue, suggesting that they could be pathogenic, and Met13, independently flagged by Level 10 as a core ligand-pocket residue, reappeared here with a direct ligand contact. Variant convergence (Level 14) provides more detailed information for each variant, among which the overlap with structural features and the functional impact in ClinVar and Alpha Missense, to facilitate prioritization.

### Accessing Atlantis through a LLMs via the MCP server: c-Src’s PTM crosstalk as a case study

The Atlantis MCP server enables users to query its human proteome-wide knowledge based on natural language prompting via an AI assistant. Once connected to the MCP server, users can interact with the Atlantis database through their preferred LLM without having to formulate API or GraphQL queries themselves. The assistant autonomously identifies the appropriate Atlantis tools and constructs the queries required to retrieve the requested information (see Methods for MCP server configuration and access). Users can ask questions such as “Which PTMs are spatially clustered at the c-Src kinase?” and retrieve the corresponding results directly through the conversational interface (Figure 5A). These results can include spatially close PTMs, their distances, and the experimental structures in which the associations are observed. The MCP interface therefore provides conversational access to the residue-level structural and functional information available in Atlantis.

**Figure 5.**
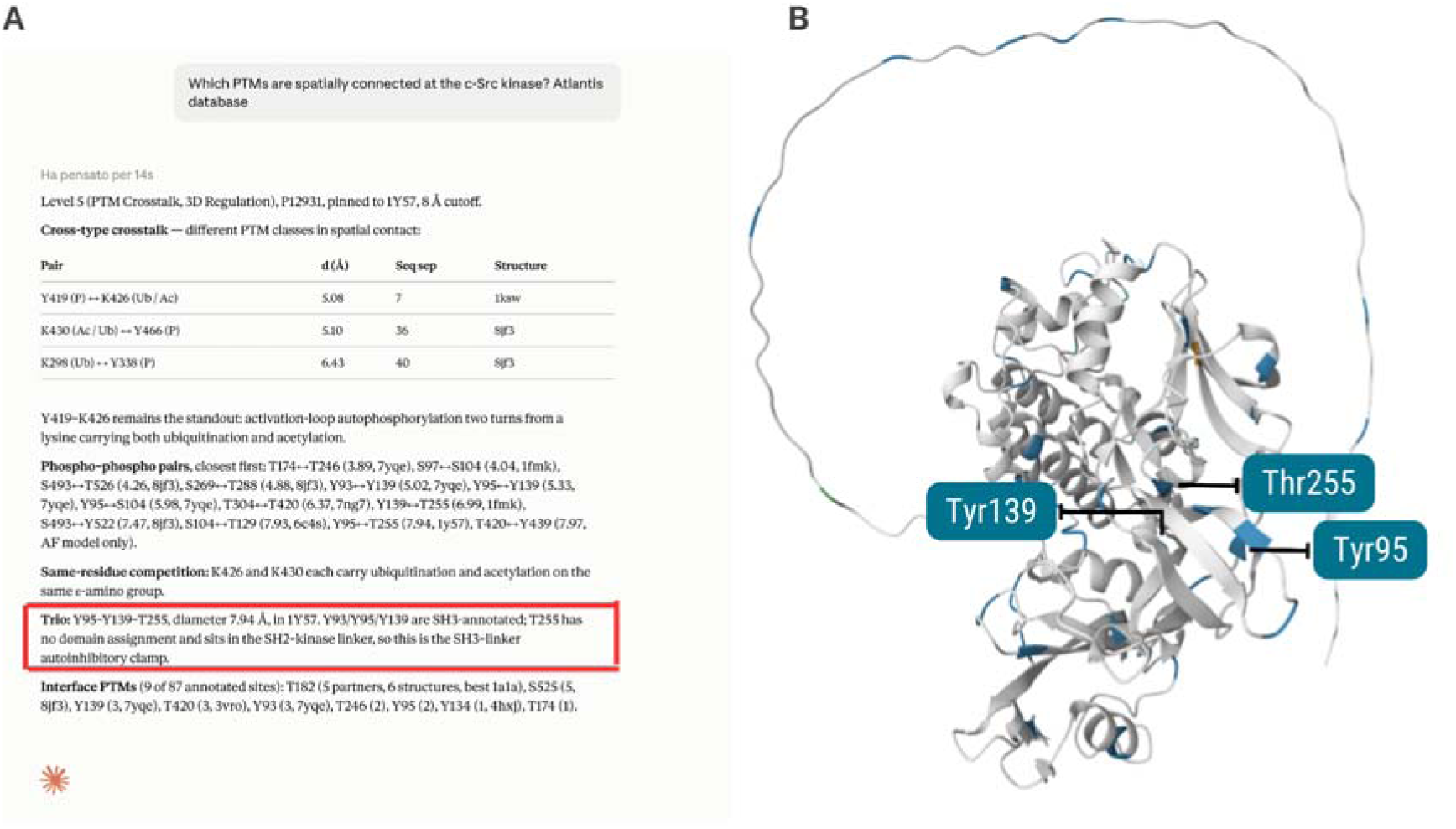
Pathfinder Level 5 (3D PTM Crosstalk) applied to c-Src kinase (UniProtKB P12931). **(A)** Natural-language query answered by an LLM agent through the Atlantis MCP server, which ran Level 5 on P12931 and returned cross-type, phospho–phospho, same-residue, and interface crosstalk sets. The three-phosphosite trio at the SH3/SH2–kinase interface is highlighted in red. **(B)** The same trio displayed in the Functional Site Information section of the P12931 protein page in the Atlantis web interface, where annotated sites are mapped onto the AlphaFold model AF-P12931 in the embedded Mol* viewer. PTM sites are shown in blue on the white cartoon, and the three clustered phosphorylation sites, Tyr95, Tyr139, and Thr255, are labelled.

c-Src is activated by conformational changes driven by its phosphorylation state, switching between a closed, autoinhibited form and an open, catalytically competent one according to the phosphorylation status of two regulatory tyrosines with opposing effects: phosphorylation of the C-terminal tail (Tyr530) stabilizes the inactive state, while phosphorylation of the activation loop (Tyr419) stabilizes the active one ^49^. Src’s activity is governed by two nested clamps: the docking of phosphorylated Tyr530 into the SH2 domain, and a second clamp in which the SH3 domain engages the SH2-kinase linker, which together lock the kinase in an autoinhibited state ^50^. Level 5 (PTM Crosstalk, 3D Regulation), which finds PTM sites in spatial proximity across intra-chain contacts, was run through an LLM agent via the Atlantis MCP server and identified a PTM cluster corresponding to this second clamp: three phosphosites (Tyr95, Tyr139, Thr255) spanning the SH3 domain and linker cluster with diameter 7.94 Å (Figure 5B). Level 1 confirmed that Tyr95 and Thr255 are in direct contact in the active-like structure (PDB 1Y57; 7.94 Å) but require a Tyr139 detour in the canonical closed structure (PDB 1FMK; 12.78 Å) (Figure 5B).

## DISCUSSION

The Atlantis database is a comprehensive platform for exploring residue-level protein information. It builds upon data provided by public resources such as UniProtKB^1^ for sequence features, the PDB^19^ and AlphaFold DB^3^ for structural context, interaction databases like IntAct^25^, PTM repositories like PhosphoSitePlus^26^, and variant collections like ClinVar^20^. While various tools allow visualization of some features on structures^2,3^ or calculation of contacts^5^, Atlantis systematically integrates these diverse data sources. Available features include structural annotations, protein architecture details, pre-calculated intra- and inter-protein interactions from both experimental and predicted structures, ligand binding information, and clinically relevant variants with pathogenicity scores, into a unified, residue-centric framework. This integrated approach helps researchers investigate the functional role of specific residues and their involvement in biological processes and diseases. The user-friendly interface and GraphQL API facilitate efficient data retrieval and integration into automated workflows, which could be used for data analysis and machine learning pipelines, providing a valuable tool for both experimental and computational biologists. The MCP connector also allows to interrogate Atlantis through LLMs, offering easier access via natural language prompting as well as new means for data integration and analysis.

Beyond data integration, Atlantis offers active analytical capabilities through its Pathfinder and Contact Calculation modules. The Pathfinder module allows the analysis of residue interaction networks and integrates it with the different types of functional annotations available in the database. For example, it is possible to inspect shortest communication paths in 3D structures, prioritize allosteric structural hubs, and mechanistically contextualize the microenvironment of specific variants or PTMs. Furthermore, the Contact Calculation module extends this powerful analytical framework to user-inputted structures. By allowing researchers to upload custom experimental models or newly predicted complexes, the platform can calculate residue contact networks and automatically map Atlantis’s extensive functional repository onto novel structures.

While Atlantis provides a robust foundation for residue-level protein analysis, there are potential avenues for further development. The current architecture is scalable and adaptable for incorporating proteomes from other model organisms. Moreover, the Atlantis database is expected to be an ideal avenue to integrate the contact analysis of AF-predicted interactomes (e.g. ^27,29^). To date, only AlphaSync^6^ provides the possibility to integrate AFDB with newly, AF- predicted structures, although multimeric predictions seem not to be currently supported.

Although some of the functionalities presented in Atlantis are available in other resources, the database’s residue-centric integrated data, comprising pre-calculated contact information from both experimental and predicted structures, and integrated ClinVar/AlphaMissense annotation, provides a comprehensive toolkit for researchers interested in gathering information about single positions in the human proteome and analyzing sets of mutations or interaction interfaces. Atlantis will serve as a valuable resource for the scientific community, facilitating the retrieval of information about protein function, molecular interactions, post-translational modifications and the impact of genetic variations. The availability of an MCP connector will streamline the integration and interrogation of Atlantis through LLMs as well as agentic frameworks for biomedical research such as Biomni ^51^.

## DATA AVAILABILITY

Atlantis webserver is freely available at: https://atlantis.bioinfolab.sns.it.

The underlying code is freely available at: https://github.com/raimondilab/atlantis_db.

## AUTHORS CONTRIBUTION

N.D.O.R. realized the webserver, performed the analysis, wrote the manuscript; P.F. performed the analysis; M.V. contributed the analysis; M.M. contributed the analysis; M.R. contributed the analysis; P.M. supervised the project, performed the analysis, wrote the manuscript; F.R. conceptualization, supervised the project, wrote the manuscript.

## ACKNOWLEDGEMENTS

We gratefully acknowledge the CINECA award, in collaboration with AIRC, for the availability of high-performance computing resources and generous support, as well as the computational resources of the Center for High-Performance Computing (CHPC) at Scuola Normale Superiore.

